# Digital embryos – A novel technical approach to investigate perceptual categorization in pigeons (*Columba livia*) using machine learning

**DOI:** 10.1101/2021.05.31.446401

**Authors:** Roland Pusch, Julian Packheiser, Charlotte Koenen, Fabrizio Iovine, Onur Güntürkün

**Author notes:** Co-first authors (Roland Pusch, Julian Packheiser).

## Abstract

Pigeons are classic model animals to study perceptual category learning. A theoretical understanding of the cognitive mechanisms of categorization requires a careful consideration of the employed stimulus material. Optimally, stimuli should not consist of real-world objects that might be associated with prior experience. The number of exemplars should be theoretically infinite and easy to produce. In addition, the experimenter should have the freedom to produce 2D- and 3D-versions of the stimuli and, finally, the stimulus set should provide the opportunity to identify the diagnostic elements that the animals use. To this end, we used the approach of “virtual phylogenesis” of “digital embryos” to produce two stimulus sets of objects that meet these criteria. In our experiment pigeons learned to categorize these stimuli in a forced-choice procedure. In addition, we used peck tracking to identify where on the stimulus the animals pecked to signal their choice. Pigeons learned the task and transferred successfully to novel exemplars. Using a k-nearest neighbor classifier, we were able to predict the presented stimulus class based on pecking location indicating that pecks are related to features of interest. We further identified potential strategies of the pigeons through this approach, namely that they were either learning one or two categories to discriminate between stimulus classes. These strategies remained stable during category transfer, but differed between individuals indicating that categorization learning is not limited to a single learning strategy.

## Introduction

Categorization can be defined as the ability of an organism to classify stimuli by responding equivalently to members of the same class, unequally to members of different classes, and to transfer this acquired knowledge to novel instances of these categories (Cohen & Lefebvre, 2017; Keller & Schoenfeld, 1950). In comparative research, pigeons have been proved to be a valuable model organism to study category learning as they have been shown to be able to categorize extremely complex visual stimuli. For examples, they can reliably categorize photographs with humans vs. those without (Aust & Huber, 2001, 2002; Herrnstein & Loveland, 1964), words vs. nonwords (Scarf et al., 2016), benign vs. cancerous histological samples (Levenson et al., 2015), and pictures healthy and diseased heart muscle (Navarro et al., 2020). Especially for these complex naturalistic categories the question arises how this well-documented ability is realized.

Different approaches have been employed to isolate diagnostic perceptual features of stimuli that drive perceptual categorization learning. One approach controls the feature content by using carefully constructed artificial stimuli to investigate the underlying categorization mechanisms. A nice example of this approach is the study of Jitsumori (Jitsumori, 1996), in which pigeons were tested with artificial butterfly figures that differed along three dimensions (darkness of shading, number of blobs on forewings, number of blobs on hindwings). These kind of experiments successfully showed that color, size, shape and their combined configural cues all seem to be behaviorally relevant (Wasserman & Biederman, 2012). However, due to the narrow detail in artificial, geometric stimuli, the problem of limited generalizability to natural visual classes remains (Lazareva & Wasserman, 2017). In addition, pigeons often encountered severe problems in mastering these tasks, which is why Lea, Wills & Ryan (Lea et al., 2006) concluded that artificial polymorphous categories do not seem to represent a good model for natural perceptual categories in birds.

Other attempts use complex natural stimuli to study animal categorization abilities (e.g. photographs of natural scenes, Fersen & Lea, 1990) at the expense of being able to clearly demonstrate the exact features the animals were responding to (Aust & Huber, 2001). The reason is possibly that in natural categories different reward predicting features co-occur. For example in a human present vs human absent categorization, humans are often depicted alongside “man-made” objects like cars, streets, furniture etc. and indeed it has been shown that pigeons can learn to categorize such “man-made objects” (Lubow, 1974). In addition, many of such features may contribute to the class assignment such that pigeons can ignore some and simply attend to few that have the highest predictive value (Huber et al., 2000). A further difficulty of these approaches is prior experience. Lab animals typically have a significant amount of experience interacting with humans and observing other objects around the lab. This turns out to be relevant in categorization tasks as pigeons with no experience seeing human heads fail to show a preference for heads over arbitrary skin patches in a categorization task (Aust & Huber, 2010). Thus, there are important limits to manipulating and altering given features when working with complex photographs because the precise identification of pictorial features exerting control over behavior is astonishingly difficult (e.g., Aust & Huber, 2002).

An approach to directly reveal diagnostic features for categorization is the bubbles procedure (Gosselin & Schyns, 2001). Using this method, randomly distributed occlusions (bubbles) obstruct the view of parts of the stimuli that are to be categorized. Based on the categorization performance, the relative contribution of the different image features can be reconstructed (Gibson et al., 2005; Gibson et al., 2007). Although this approach represents an advance, large parts of the stimuli are occluded in test trials. Thus, the animals are forced to use local processing strategies as global strategies are prevented by occlusion.

A novel approach to address these shortcomings is to introduce “digital embryos” (see figure 1). This novel stimulus type was created by Hegdé and colleagues as a versatile tool for categorization learning research (Hauffen et al., 2012; Hegdé et al., 2008; Kromrey et al., 2010). Both humans (Hegdé et al., 2008) and macaques (Kromrey et al., 2010) can learn to categorize those embryos and are able to transfer their knowledge to novel instances. Digital embryos are developed from a parent object using “virtual phylogenesis” (VP); a process that emulates biological evolution that, in this case, is under human guidance. The evolutionary computer algorithm can create any number of classes with varying degrees of complexity and difficulty in discrimination. Thus, digital embryos mimic natural objects and possess features that are also readily controlled by the experimenter. Employing these artificial and yet naturalistic stimuli enables us to study perceptual categorization without having to worry about prior knowledge of the stimulus set.

**Figure 1.**
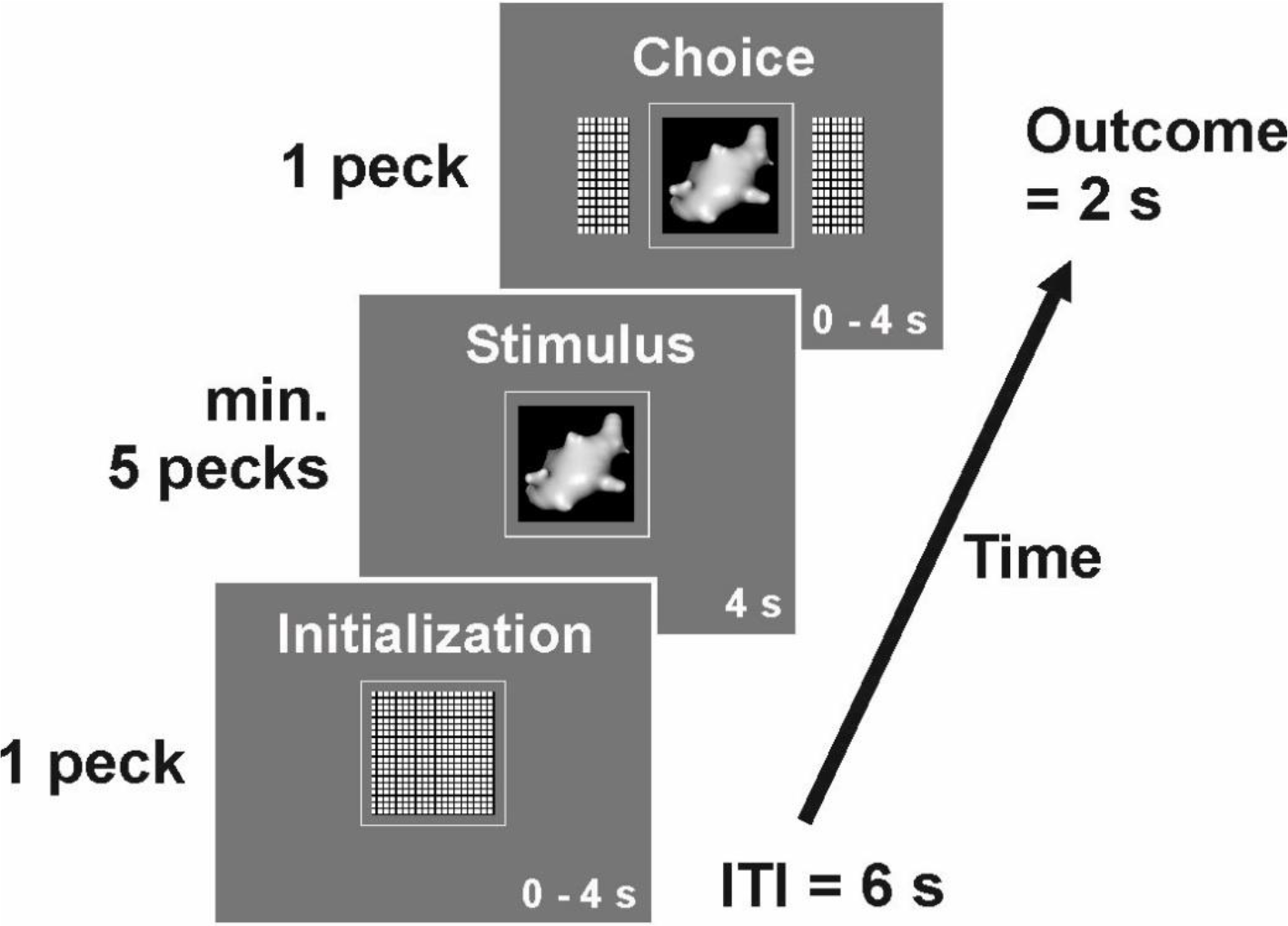
Example trial of the categorization task. Each trial began with a short high tone. First, the initialization key was presented for up to 4 s and could be terminated once pecked. Then, one digital embryo stimulus was presented (depicted is a digital embryo out of class X) and had to be pecked at least 5 times. It could not be terminated for a fixed time interval of 4 s. Subsequently, two choice keys were presented alongside the stimulus for 4 s. One peck on either key indicated the pigeon’s choice and was either immediately rewarded with 2 s access to grain or punished with lights turned off and a low punishment tone for 2 s. For half of the pigeons, the right choice key represented class X and the left choice key represented class Y (vice versa for the other half of the pigeons). The next trial followed after a 6 s inter-trial interval.

A method to identify the informative diagnostic variables of a specific category that pigeons utilize in a categorization task is to directly reveal the focus of attention during categorization. To this end, peck tracking was introduced to identify the pecking locations of pigeons (Allan, 1993; Dittrich et al., 2010; Jitsumori & Yoshihara, 1997). Dittrich et al. (2010) could indeed show that pigeons working on a people present/people absent categorization task preferentially pecked on the heads of the depicted persons and were subsequently impaired in their performance when they had to run the task with the heads of the humans removed from the photographs. Similarly, Castro & Wasserman (2014, 2017) also demonstrated that pigeons track features that are category-relevant and respond less to details that only weakly coincide with the presence of the relevant stimulus. This patern of behavior would be predicted from the “Common Elements” theory of categorization that assumes that images from a category contain some common features (Soto & Wasserman, 2010). Choosing pictures with such features correlates with reward probability and consequently drives learning to specifically attend to such common elements.

The present study combines the peck tracking method with the digital embryo approach to study which stimulus features are used for object categorizations. By employing a machine learning approach we aim to identify whether pecking locations were predictive of the presented category and to reveal the underlying learning strategy of individual animals. In our experiments, we trained pigeons to distinguish between two different classes of digital embryos in a single interval forced-choice task. First, the animals went through a category learning procedure and second, conducted a transfer test with new stimuli of the same class. Peck tracking using touch-screen technology allowed us to gain insight into local attentional mechanisms and categorization strategies. Our hypotheses of the following experiments were: (*i*) Pigeons, in line with humans and monkeys, can discriminate between the two classes of digital embryos. (*ii*) Categorical information can be transferred to new exemplars of a given category. This can be viewed as a proof of open-ended categorization, a level beyond rote categorization (Herrnstein, 1990). (*iii*) Using machine learning, we can successfully predict the presented stimulus class only based on pecking location ultimately providing insight into the features used for discrimination as well as the underlying strategy of each individual animal.

## Methods

### Subjects

Eight unsexed adult homing pigeons (*Columba livia*) obtained from local breeders served as subjects. The pigeons were housed in individual wire-mesh cages with a 12-hour light-dark cycle beginning at 8:00 a.m. They had free access to water and were food deprived, maintained at approximately 80-90 % of their free feeding body weight and fed accordingly with a mixture of different grains. The subjects were treated in accordance with the German guidelines for the care and use of animals in science. All procedures were approved by a national ethics committee of the State of North Rhine-Westphalia, Germany. All experimental conduct was in agreement with the European Communities Council Directive 86/609/EEC concerning the care and use of animals for experimental purposes.

### Apparatus

The experiment was conducted in a custom-built operant chamber (35 × 35 × 35 cm) with a touch-screen monitor (model ET1515L with APR technology, Elo Touch Solutions Inc., Milpitas, CA, USA), on which three horizontally aligned areas were defined as pecking keys (cf. figure 1; central pecking key 5 × 5 cm, lateral pecking keys 3.5 × 5 cm with a 1 cm gap in between pecking keys; 17 cm above the chamber floor).

The touch-screen was used to present the stimuli, record pecks and location coordinates for each peck. The stimuli measured 4 × 4 cm and were presented within the central pecking key, surrounded by a 0.5 cm gray border. The lateral pecking keys measured 2.5 × 4 cm, surrounded by a 0.5 cm gray border that blended with the gray background of the touch-screen. In both cases, pecking this gray border was registered as a valid peck. This buffer was included because there is no distinct physical pecking key on touch screens and pigeons sometimes peck on the edge of a stimulus display. The grey border enables these pecks to be included. A centrally located food hopper delivered mixed grains as reward. Experimental hardware was controlled with custom-written Matlab code using the Biopsychology-Toolbox (Rose et al., 2008).

### Stimuli

Two classes of digital embryos, arbitrarily termed X and Y, were generated using the software Digital Embryo Workshop (Brady & Kersten, 2003; Hauffen et al., 2012). These classes were created using a process called “virtual phylogenesis” an algorithm that mimics biological evolution by creating multiple virtual taxonomic classes of 3D objects that have been termed “digital embryos”. This process starts with an icosahedron (a polyhedron with 20 faces) as the parent object. Over several virtual generations multiple separate classes “evolve”, each of which shares identifying within-class characteristics. The system allows the production of an endless number of individuals that are members of one or the other class. Example embryos from different generations including class X and Y are depicted in figure 2 and full sets of stimuli are depicted in supplementary figures 1 and 2. In our study, class X and class Y consisted of 30 members each, 20 of which were randomly chosen for category training and 10 used for transfer tests.

**Figure 2.**
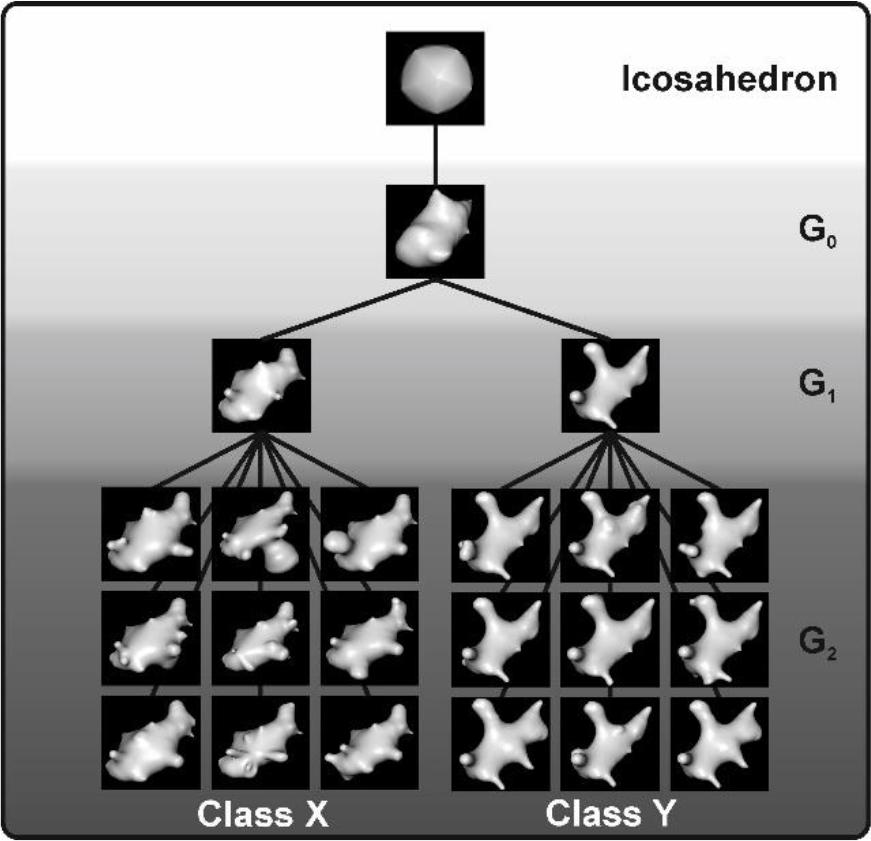
Digital embryo classes. Digital embryos were created with a process called “virtual phylogenesis” that mimics biological evolution. Starting with an icosahedron, different generations (G_0_-G_2_) are created, resulting in embryo classes X and Y.

### Behavioral Task

Pigeons were trained in daily sessions (5 days a week) on a single interval forced-choice task (figure 1). The start of each trial was indicated by a short high frequency tone. Pigeons first had to peck at least once on an initialization key in a 4 s time window to commence the trial. Then, one digital embryo stimulus was presented for a fixed time interval of 4 s. During this period the pigeons were required to peck at least 5 times on the stimulus. If they did not, the trial was aborted. If the bird had pecked five times during the 4 s interval, two choice keys were presented alongside the stimulus. Subsequently, pigeons had to make a choice, i.e. peck once on either the left or right choice key within a 4 s time window. The stimulus-response association was balanced across pigeons. When a choice was made, the display disappeared. For correct choices, the animals had 2 s access to food and the feeding light was turned on. For incorrect choices, the ambient chamber lights were turned off for 2 s accompanied by a low frequency tone. The next trial followed after a 6 s inter-trial interval (ITI). For experimental reasons, the paradigm had to be divided into two phases.

#### 1: Embryo Learning Phase

During an initial training phase, pigeons learned to peck on all three pecking keys in a standard autoshaping procedure. Subsequently, they learned to peck the central key first, and then one choice key second, in order to obtain a food reward. During category training, all stimuli (20 class X, 20 class Y) were presented 5-10 times in a pseudo-randomized order. Eighty percent of trials were single-choice trials, i.e. only the correct choice key was visible. The remaining 20 % of the trials were free-choice trials. This means both choice keys were visible, could be pecked on, and errors could be made. Single-choice trials were gradually reduced to zero once pigeons correctly categorized the stimuli in more than 80 % of the free-choice trials in two consecutive sessions. Additionally, the probability of food reward was gradually reduced from 100 % to 60 % to ensure stable behavioral performance in longer sessions.

#### 2. Transfer Test Phase

In the transfer test phase, pigeons’ choices to transfer stimuli, i.e. new whole embryo exemplars of class X and Y and choices to known stimuli, i.e. embryo exemplars learned during training, were tested. The stimulus sequence consisted of 20 known whole embryos of each class, which were repeatedly shown 4 times (80 class X + 80 class Y = 160). These trials are labeled known-stimulus trials. Further, the sequence included the presentation of 10 new embryos (10 class X + 10 class Y = 20). These trials are labeled transfer-stimulus trials. Each individual session comprised up to two of these blocks, with a reward probability ranging between 43 – 72 % (mean: 57 %) depending on the individual performance of the animal.

In a first transfer test phase, animals received no feedback for behavioral responses in transfer-stimulus trials, thus, reward or punishment was omitted irrespective of the animals’ responses. In this manner, transfer stimuli could be tested without reward contingencies influencing the pigeons’ decision. Unfortunately, under these conditions, four pigeons showed a neophobic response and refused to peck altogether. Therefore, we repeated the transfer phase during which all pigeons received non-differential reinforcement for transfer trials, i.e. food reward irrespective of correct or incorrect choice (van Hamme et al., 1992). This motivated the animals to attend to the task without differentially reinforcing one response over another in test conditions. We will only report the results from the reinforced transfer test in the main results as the behavioral performance during the non-reinforced and reinforced transfer were virtually identical for the four pigeons who responded in both transfer tests (results for the non-reinforced transfer can be found in the supplementary data).

### Data Analysis

Pigeons’ pecking and choice behavior was stored and subsequently analyzed in MATLAB Version 2020a. We calculated percentage of correct choices collapsed for both classes X and Y for each animal individually. This was done for known and transfer stimuli across both experimental phases. We then calculated one-sample t-tests to identify if performance was different from chance-level (50% as the choice was binary). To identify if performance on transfer trials differed from performance on known-stimuli trials, a paired t-test was conducted. The same procedure was repeated for stimuli of class X and Y separately. Furthermore, we used precise peck tracking to gain insight into the pigeons’ center of attention. Using touch-screen technology x- and y-coordinates of each peck in the stimulus/border area were stored and analyzed for each stimulus presentation in both known- and transfer-stimulus trials. If the pigeon pecked outside the stimulus/border area, data on these pecks was discarded. Pecks on all stimuli of class X and Y, respectively, were collapsed and displayed in heat-maps separately for each pigeon to visualize each animal’s center of attention. To create heat-maps, the image was sectioned into 15 × 15 equally sized squares (figure 3). Each single square covered an area of 0.11 cm^2^ of the stimulus display. Pecks located in each square were summed and because the total number of pecks differed between pigeons, the relative amount of pecks (relPecks = (100 / allPecks) * pecks/square) is depicted in the heat-maps.

**Figure 3.**
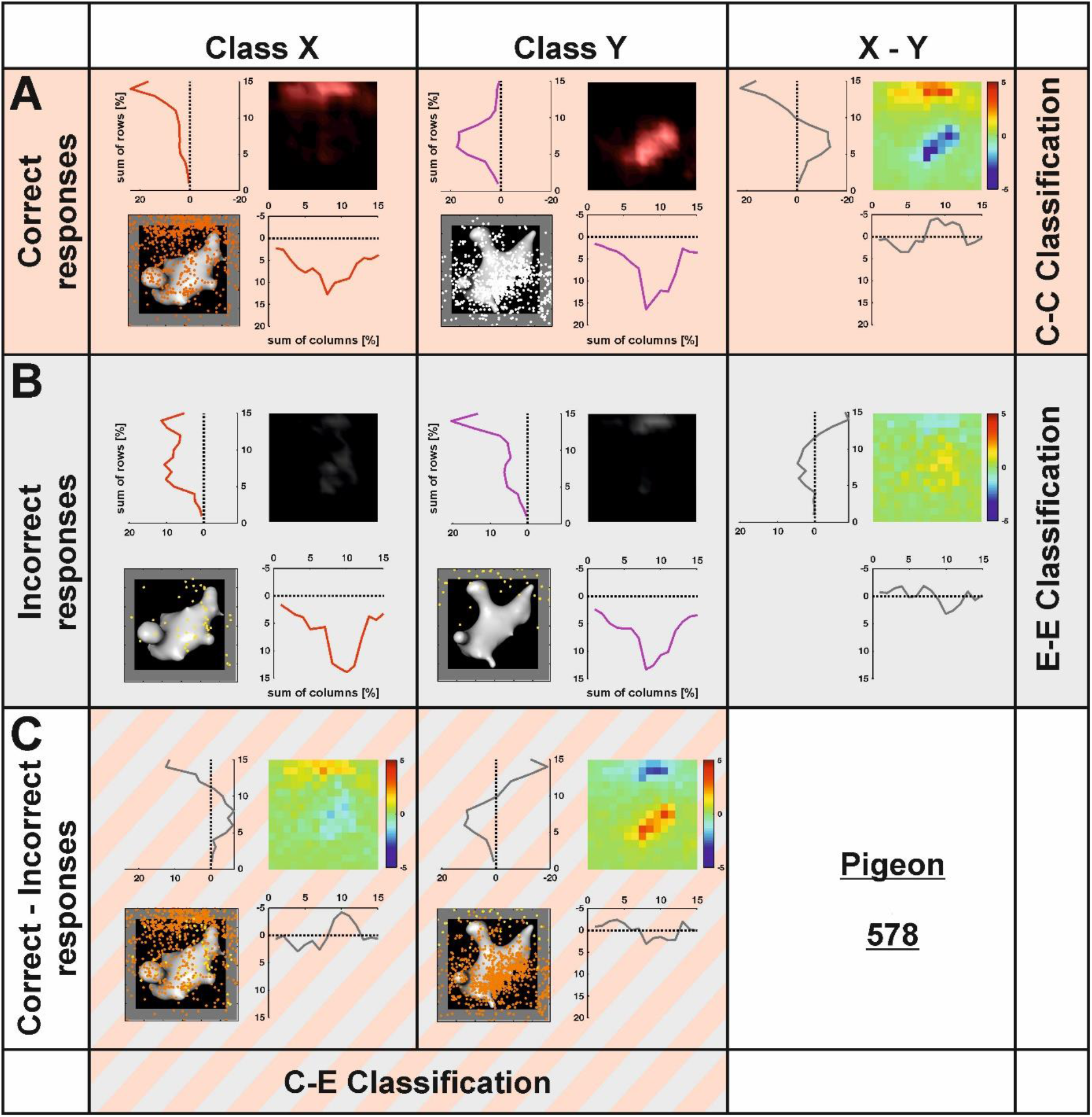
Rationale of the heatmap analysis exemplified for pigeon 578. In row **A,** the peck distribution for correctly responded stimulus presentations is shown separately for stimulus class X and Y. To create heatmaps, the stimulus display was divided in sections of 15 × 15 squares and pecks were counted in each section. Heatmaps were created based on relative pecks to compare individual animals. The subtraction of both heatmaps reveals the different locations of pecks for each stimulus class. The precise location of each peck is used by the classifier to separate both classes resulting in the Correct-Correct (CC) classification. Row **B** shows the peck distribution for incorrectly responded stimulus presentations. The subtraction of both heatmaps reveals the different locations of pecks for each stimulus class in error trials. The precise location of each peck is used by the classifier to separate both classes resulting in the Error-Error (EE) classification. Row **C** depicts the subtraction of correct and incorrect heatmaps for each class separately resulting in the Correct-Error (CE) classification. The depicted analysis is exemplary based on the responses of pigeon 578. Response profiles of each individual pigeon is given in supplementary figures 3 - 9.

### Classifier analysis

To identify if peck locations were behaviorally relevant for each individual animal, we used a k-Nearest Neighbor (kNN) classifier algorithm to predict the presented stimulus class based on the pecking location. The classifier analysis was performed in known- and transfer-stimuli trials. For all correctly responded trials, we randomly sampled n = 250 pecking events from the behavioral session and trained the algorithm as follows: the x and y coordinates of each peck were associated with the associated label (0 for class X trials, 1 for class Y trials). A classification was then computed using k = 15 nearest neighbors according to the recommendation to use an uneven number of neighbors as well as k = sqrt(n) of training events for good classification accuracy and low computational expenses (Lall & Sharma, 1996).

In a next step, 250 randomly chosen pecking events from correct trials (different from the training events) were labeled according to the kNN classifier based on their pecking location. Accuracy of the classifier was measured by quantifying the success rate of correctly classifying the presented stimulus class. Classification was performed across 10 iterations each feeding random pecking events as training trials and predicting another subset of test trials to cross-validate the input and output of the algorithm and identify the variance in accuracy. To quantify if the results from the classifier were statistically significant from a chance-based estimate, we repeated 10 iterations of the classifier, but fed the algorithm with shuffled input where class labels and x- and y-coordinates were permutated randomly. A dependent t-test between the results from the empirical and shuffled data was then performed. Since this analysis trained and tested the algorithm with pecking events from correct trials, we will refer to this classification as Correct-Correct or CC classification.

If sufficient pecking events occurred after which the animals made erroneous choices (> 500 events), we also quantified if pecking locations from correct trials were predictive of error trials to identify whether pecking in correct trials and error trials was consistent or differed from one another. For that purpose, the aforementioned analysis was repeated, but test pecks for the algorithm were chosen from erroneous trials rather than correct trials. This analysis will subsequently be referred to as Correct-Error or CE classification. Furthermore, if sufficient pecking events from error trials occurred, we also predicted the class label using error trials as both training and test events. Aim of this analysis was to quantify the internal consistency in pecking during error trials, i.e. if the stimulus class in error trials can be predicted based on pecking location in error trials. This analysis will be dubbed Error-Error or EE classification.

## Results

### 1. Embryo Learning Phase

The embryo learning phase took between 29 and 49 sessions to meet criteria and move on to the next phase. All eight pigeons performed two consecutive sessions at 80 % correct of the free choice trials before moving to the next phase. After reaching the initial learning criterion, a step-wise reduction of single-choice trials as well as of reward probability followed to move to the transfer test phase.

### 2. Transfer Test Phase

All eight pigeons were successfully tested on known- and transfer-stimuli during the test session when transfer trials were rewarded non-differentially. Overall, performance in known-stimuli and transfer-stimuli trials was very high (Mean = 91.81 %, SD = 3.26 %; Mean = 91.36 %, SD = 3.80 %, respectively, see figure 4) and was significantly different from chance (t(7) = 4260.45, *p* < .001, Cohen’s d = 12.83; t(7) = 3.654.37, *p* < .001, Cohen’s d = 10.89, respectively). There was no difference in performance between known- and transfer-stimuli trials (t(7) = 0.61, *p* = > .250, Cohen’s d = 0.45). Individually, the two stimulus classes, both class X (known-stimuli trials mean = 92.89 %, SD = 3.23%; transfer-stimuli trials mean = 92.58 %, SD = 3.03 %) and class Y (known-stimuli trials mean = 91.26 %, SD = 3.98 %; transfer-stimuli trials mean = 90.13 %, SD = 6.55 %), showed above chance-level performance (all *p*s < .001). There was no difference between stimuli of class X and Y in both known-(t(7) = 0.97, *p* > .250, Cohen’s d = 0.32) and transfer-stimuli trials (t(7) = 1.02, *p* = > .250, Cohen’s d = 0.42). Behavioral results of the four pigeons who completed the transfer test in the absence of reward are presented in the supplementary results.

**Figure 4.**
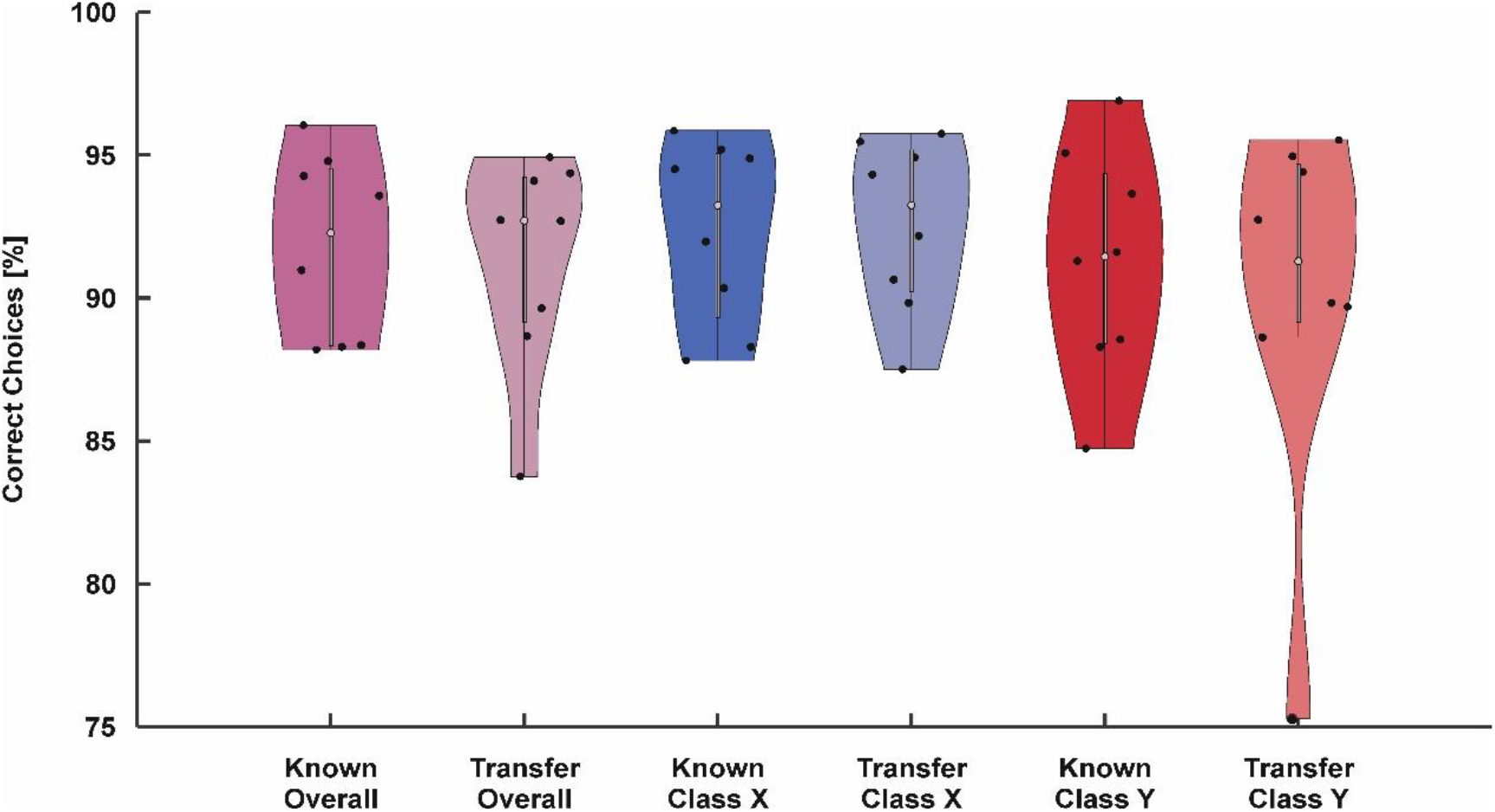
Behavioral results. Violin plot of behavioral performance for known and transfer trials overall and broken down by embryo class. Digital embryos could be categorized in each experimental condition under non-differential reward conditions. There was no difference between the performance to the known embryos and the transfer to new instances of embryo classes X and Y in any of both conditions.

### 3. Peck Tracking

We used a k-nearest neighbor classifier to identify whether the presented stimulus class can be predicted based on the pecking location. To this end, the classifier was trained with randomly chosen pecking events across behavioral sessions in which pecking events were labeled according to the presented stimulus class in a given trial. Another subset of trials was then used to determine the predictive accuracy of the classifier based on pecking location alone (see Methods). This was done for each animal separately. Results of all analyses for the four pigeons that successfully performed the non-reinforced transfer are presented in the supplementary data.

Our aim was to investigate whether the class label can be predicted in Correct-Correct or CC classification in which both training and test trials were constituted by correct trials. All eight animals could be investigated for a CC classification in known-stimuli and transfer-stimuli trials. For known-stimuli trials, we found that the classifier could significantly predict the stimulus class based on pecking location for seven out of eight pigeons (P578: 82.08 %, P580: 80.84 %, P582: 70.56 %, P583: 64.24 %, P592: 77.84 %, P593: 64.64 %, P599: 77.80 %, all *p*s < .001). Only results for pigeon P579 did not differ significantly from chance (52.40%, t(9) = 1.22, *p* > .250, Cohen’s d = 0.29, see figure 5A) as the animal indifferently pecked on one side of the stimulus irrespective of the presented stimulus class. Animals preferred different embryo features indicated by their idiosyncratic pecking locations for the two stimulus classes (figure 6).

**Figure 5.**
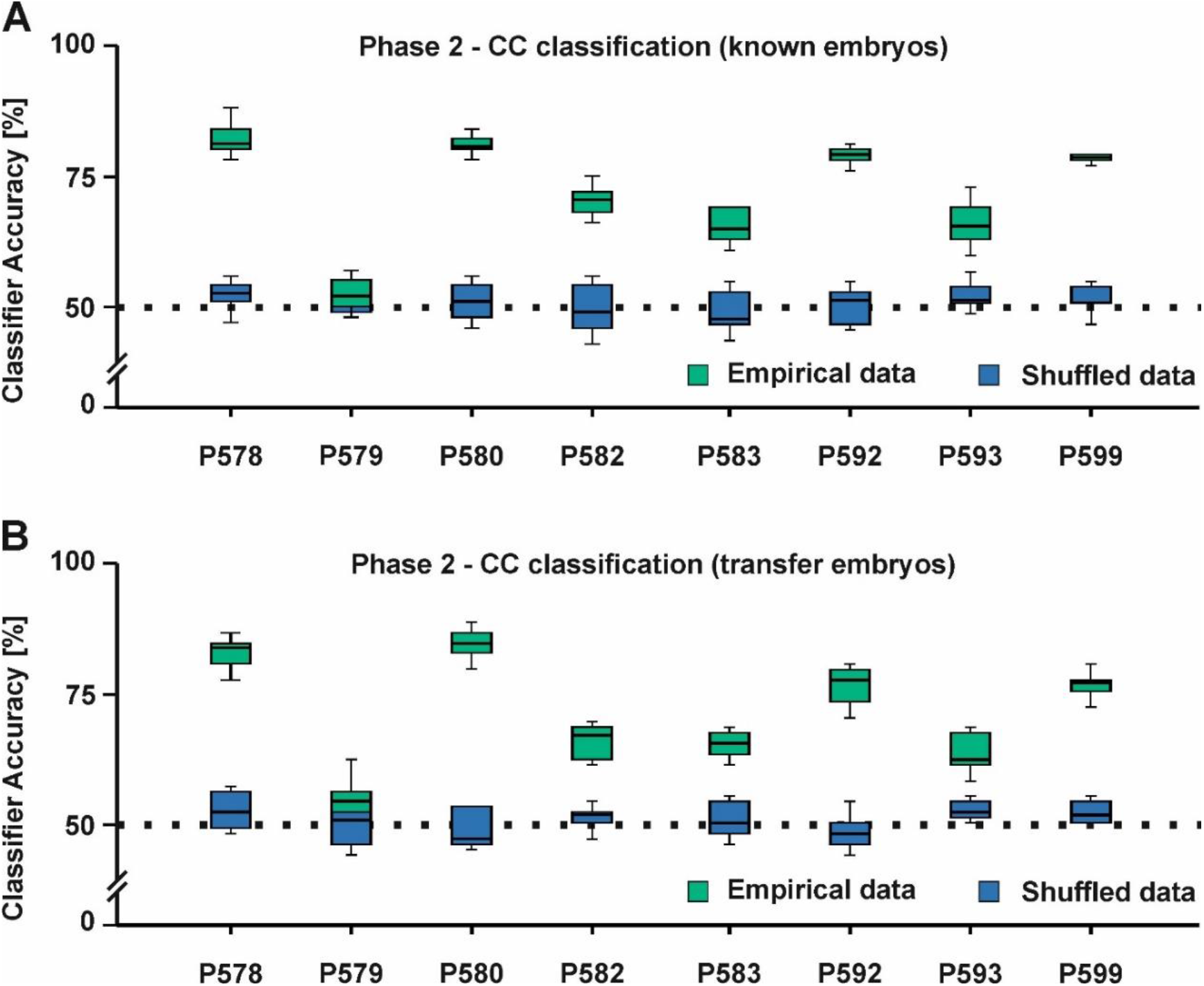
CC classification results of the second experimental stage. Digital embryos could be classified for each animal tested (**A** shows the classifier response for the known stimuli and **B** depicts the classifier results for the transfer stimuli).

**Figure 6.**
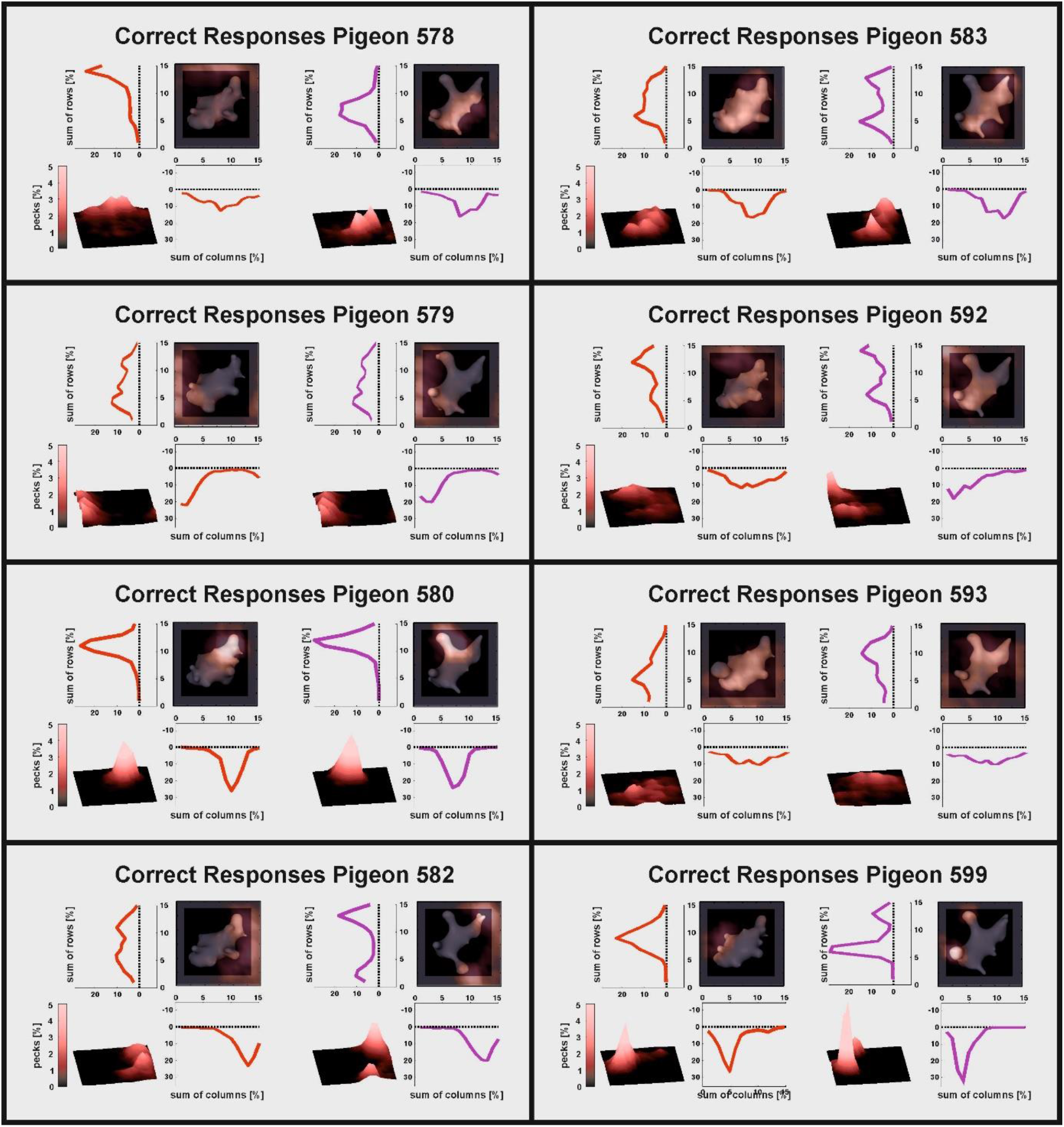
Heatmap analysis for all individual pigeons. Only pecks from correct trials for the two stimulus classes (left: X, right: Y) are shown. Additional analyses as presented in figure 3 are given in the supplementary figures.

For transfer-stimuli trials (figure 5B), we found identical results compared to known-stimuli trials (P578: 82.09 %, P579= 53.48 %, P580: 83.68 %, P582: 65.52 %, P583: 64.96 %, P592: 76.12 %, P593: 63.16 %, P599: 76.20 %, all *p*s < .001). There was no difference from known-stimuli trials in CC accuracy except for pigeon P582 for which a slightly lower classification accuracy in transfer-stimuli trials could be detected (t(9) = 2.89, *p* = .018, Cohen’s d = 0.92).

For all eight animals, there were sufficient errors allowing for a Correct-Error or CE classification in known-stimuli trials in which the classifier was trained with correct trials and tested on error trials. Here, three pigeons demonstrated similar pecking patterns in correct and error trials (indicating a within class error) as the classifier yielded above chance results in a CE classification (P580: 58.36%, P582: 62.36%, P599: 61.16%, all *p*s <.008, figure 7A). The accuracy was lower for CE compared to CC classification for all three pigeons (all *p*s < .001) indicating that pecking in error trials was more dispersed compared to correct trials.

**Figure 7.**
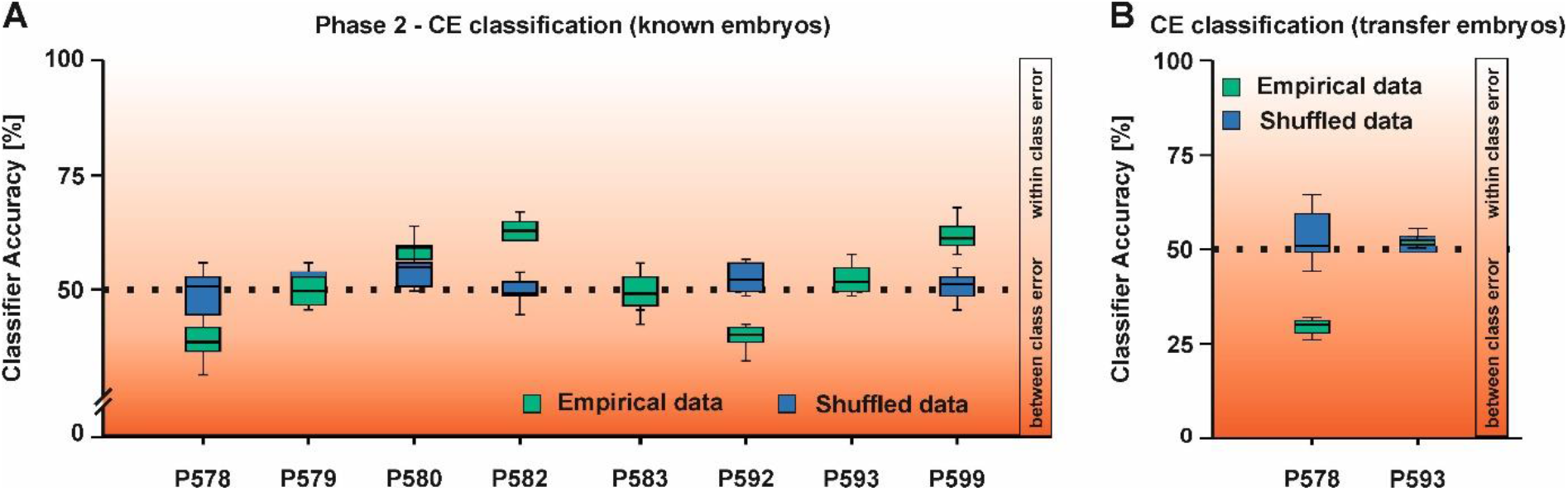
CE (Correct-Error) classification results of the second experimental stage. **A** shows CE classification for the known stimuli. **B** depicts the CE classification for the transfer stimuli. Dark red colors indicate a confusion between the categories X and Y. Light red colors indicate confusion within a given category in the CE classification.

For pigeons P578 and P592, we found that CE classification was significantly below chance (P578: 38.12 %, P592: 38.96 %, both *p*s < .001) indicating that they mixed up the stimulus classes in error trials (indicating between class errors). The last three pigeons demonstrated no significant difference from chance in the CE classification (P579: 49.16 %, P583: 49.00 %, P593: 52.24 %, all *p*s > .174) indicating that there was no relationship between correct and error trial pecking locations. For two pigeons, sufficient errors were made in transfer-stimuli trials enabling further analysis. For P578, we again found a significantly lower classification accuracy compared to chance (29.64 %, t(9) = 12.64, *p* < .001, Cohen’s d = 3.97, figure 7B). For pigeon P593, there was no association between correct and error trial pecking locations (52.70 %, t(9) = 1.55, *p* = .156, Cohen’s d = 0.49). Both pigeons thus showed consistent patterns in known-stimuli and transfer-stimuli trials.

Error-Error or EE classification in which the classifier was both trained and tested on pecking events during erroneous trials could be performed for all animals in known-stimuli trials. Aim of the analysis was to determine whether pecking in error trials was consistent. Classifier accuracy was above chance for six of the eight animals (P578: 54.64 %, P580: 63.16 %, P582: 61.64 %, P583: 74.44 %, P592: 57.52 %, P599: 72.96 %, all ps < .025) indicating that pecking locations in error trials were to a certain degree consistent rather than random. For pigeons P579 and P593, no significant difference from chance could be found (both ps > .270). For pigeons P578 and P593, there were sufficient error events in transfer-stimuli trials to allow for EE classification. Here, results were comparable to known-stimuli trials in these pigeons as classification accuracy was above chance for pigeon P578 (67.6 %, t(9) = 5.95, p < .001), but at chance-level for pigeon P593 (53.92 %, t(9) = 1.29, p = .228). The EE classification accuracy was however significantly higher in transfer- compared to known-stimuli trials (t(9) = 10.21, p < .001)

## Discussion

In the present study, we investigated perceptual categorization learning in pigeons using “digital embryos” as visual stimuli. The process of “virtual phylogenesis” can generate any number of these artificial stimuli and categories while maintaining natural stimulus characteristics. We hypothesized that pigeons can both *(i)* learn to categorize digital embryos of different stimulus classes as well as *(ii)* transfer their knowledge onto novel exemplars of these classes. Our results clearly demonstrate that all animals were able to discriminate between classes X and classes Y and could transfer this knowledge to never before encountered stimuli of the same class. Finally, we hypothesized that *(iii)* pecking behavior is indicative of the pigeon’s attention towards critical stimulus features and that we could therefore predict the presented stimulus class solely based on pecking locations. Indeed, we could decode the presented stimulus class for the majority of the pigeons (7 out of 8) using a k-nearest neighbor approach. Classification accuracy was highly comparable between training and transfer trials. However, classification accuracy was not uniform across animals with some pigeons showcasing high accuracy classification results whereas other pigeons showed lower decoding accuracy that was nonetheless significantly different from chance level.

Peck tracking using touch-screen technology serves similar purposes as eye tracking in primate studies. Pigeons track relevant aspects of complex visual displays in discrimination tasks by pecking on them (Castro & Wasserman, 2014, 2017; Dittrich et al., 2010). Thus, the systematic analysis of the pecking behavior can give insight into attentional mechanisms. On the descriptive level, we found that most animals favored different features of the digital embryos as there was little overlap among animals regarding most pecked area for stimulus classes if pecking is used as a proxy for attention (figure 6). This strongly suggests that pigeons rely on local stimulus features, in line with the well-documented “local precedence effect” in pigeons (Cavoto & Cook, 2001; Cerella, 1980). In addition, Yamazaki et al. (2007) found that pigeons can categorize highly scrambled pictures of humans based on small local features, thereby neglecting the overall stimulus configuration. In a comparative approach, (Aust & Braunöder, 2015) conducted both an exemplar and rule-based categorization task in pigeons and humans and found that pigeons rely on local features whereas humans have no strict preference for local or global information to solve the tasks. However, a variety of studies also found global strategy-based categorization in pigeons, indicating that they are capable of using both strategies (e.g., Goto et al., 2004; Yamazaki et al, 2007). Possible explanations for this discrepancy might be due to differences in experimental procedures and related task-demands. Goto et al. (2004) found that pigeons prefer global strategies if stimuli are relatively small resulting in a small effective visual angle during responding. The stimuli we used in our study had a size of 4 × 4 cm and were at least twice the size of the stimuli used by Goto et al. (2004, ~ 2 × 2 cm). Besides the stimulus size, Goto et al. (2004) used densely packed local information in their stimulus material. Dense packing of elements has been demonstrated to promote global preference effects in humans (Dukette & Stiles, 2001). Since our stimuli were large and did not feature dense stimulus elements, a local approach was likely to be expected.

While a descriptive analysis of pecking location can provide insight into the use of local or global preferences, it cannot illuminate on the underlying learning strategy to ultimately solve the task. Our classifier analysis indicated different strategies of the animals with regards to how many categories were likely learned during the categorization process. Since our task employed a force choice procedure, the categorization could theoretically be solved by only learning about one stimulus class. For example, if only class X had been learned, the animal simply needed to identify if the stimulus is of class X or not X to determine the upcoming choice. In the CC classification, such a learning pattern most likely corresponded to an intermediate classification accuracy (roughly between 60 % to 70 %) as the animals demonstrated a clear pecking pattern for one stimulus class in correct trials, but a rather unfocused pecking pattern for the other stimulus class. If both stimulus classes received specific pecks onto relevant class features, classification accuracy jumped to levels around 80% since the classifier was trained with information about class X and Y rather than a single class. Four pigeons seemed to therefore learn about both stimulus classes whereas three pigeons seemed to only pay attention to features of one stimulus class. Interestingly, one pigeon pecked on the left side of all stimuli, regardless of the presented stimulus class (see supplementary figure 3), but still performed the task at a very high level. Thus, there is no absolute necessity to peck onto informative features to eventually perceive and process them. Since this was however only one animal, it seems to be the exception rather than the rule.

We furthermore used the classifier to inform about why errors occur in specific animals by training it with correct trials and testing it on error trials (CE classification). We found that for three animals during the second stage of the transfer test phase, CE classification simply dropped the classifier accuracy, but it remained significantly above chance (~60 %). These animals therefore made within-category errors, i.e. they pecked onto similar features in error trials as in correct trials, but these features were less focused indicating a lack of attention (Dittrich et al., 2010). This likely led to a fail in categorization as the animals could not sufficiently extract relevant features. Two animals demonstrated inversed pecking patterns whenever they made an error as indicated by a below chance classification (~35 %). Thus, they pecked on class Y features during class X trials and vice versa. These errors would be called between-category errors as the animals mix-up the two classes. Obviously, between-category errors occurred only in animals that learned about both categories according to the CC classification. Finally, three animals showed no difference from chance-level performance in CE classification indicating that they pecked randomly during error trials possibly reflecting the failure to recognize any class-specific features.

Our results fit well into the contemporary view of categorization learning in pigeons (Güntürkün et al., 2018; Soto & Wasserman, 2010). Pigeons used several, different stimulus features to solve the categorization task. Individual differences in the selection of local stimulus features, as present in our pigeons, is in line with previous reports that used peck tracking to indicate the areas of attention in complex stimulus displays. Jitsumori & Yoshihara (1997) reported different strategies in individual pigeons when categorizing angry and happy facial expressions. Some pigeons attended more to the area of the eyes, some more to the mouth. Peck location data indicating that individual pigeons focusing on different parts of the stimulus tells us that different features are utilized within the same task. These category specific stimulus features might be discernable from neuronal population responses in visual associative areas in the pigeon brain (Azizi et al., 2019; Koenen et al., 2016). In addition, Castro et al. (2020) found in their experiments that with increasing choice accuracy, pecks directed onto relevant features of the categories increased. Thus, during learning the amount of attention to reward predicting cues increased and was clearly signaled by the pecking behavior of the animals. Based on reward contingencies associated with the resulting responses, the feature that was the best predictor of reward becomes the feature that predominantly, but not exclusively controlled the pigeon’s behavior (Castro et al., 2020). This kind of feature selection does not seem to be based on feature configuration but rather on the additive integration of individual features or common elements (Jitsumori & Yoshihara, 1997; Soto & Wasserman, 2010). This process strictly follows predictions of influential learning theories (e.g., Rescorla & Wagner, 1972) and can be explained by a dopamine-mediated reduction of associated prediction errors (Schultz, 1998) that have also been demonstrated analogously in the pigeon brain (Packheiser et al., 2021). Thus, the consistent behavioral performance of individual pigeons in categorizing digital embryos might be based on common stimulus elements and their reward prediction acquired during perceptual categorization learning.

Taken together, our study provides insights about both the center of attention in pigeons during categorization as well as the underlying learning strategy using a machine learning approach. Pigeons successfully transferred class knowledge to members of each embryo “family” or class that they haven’t seen before, using multiple strategies to do so. Generally, most animals used a local approach with the exception of one animal that did not peck on relevant features at all. Interestingly, there has been a strong record of asymmetrical lateralization in birds proclaiming that the left hemisphere is dominant in local feature extraction whereas the right hemisphere is dominant in processing global feature extraction (Yamazaki et al., 2007; Güntürkün et al., 2020). It would be interesting to investigate in future studies if the behavioral strategy could be altered by simple eye-occlusion procedures to bias the processing of a particular hemisphere in birds. Regarding the learning strategy, a number of animals used the dichotomous nature of the task by likely only learning about one category. It would be highly interesting to investigate the corresponding neural correlates of these different strategies and to see how relevant visual associative layers such as the NFL (Koenen et al., 2016) and MVL (Azizi et al., 2019) encode stimulus classes with respect to the chosen categorization strategy.

## Acknowledgements

Supported by the Deutsche Forschungsgemeinschaft through SFB874 (B5) project number 122679504 to OG and through SPP 2205 - project number 430157321 to RP. We thank Alexis Garland and Samuel Thiele for their support of this study.

## Supplementary results

In total, four pigeons conducted the first stage of the transfer test phase in which the transfer-stimuli trials did not yield any feedback such as reward or punishment. Performance in known-stimuli and transfer-stimuli was generally high (Mean = 93.49 %, SD = 4.35 %; Mean = 92.31 %, SD = 3.31 %, respectively, see supplementary figure 10) and was significantly different from chance (t(3) = 2256.53, *p* < .001, Cohen’s d = 10.00; t(3) = 2964.65, *p* < .001, Cohen’s d = 12.78, respectively). There was no difference in performance between known-stimuli and transfer-stimuli trials (t(3) = 0.83, *p* = > .250, Cohen’s d = 0.35). For the stimulus classes individually, both class X (known-stimuli trials mean = 94.08 %, SD = 5.36 %; transfer-stimuli trials mean = 91.47 %, SD = 6.07 %) and class Y (known-stimuli trials mean = 92.89 %, SD = 3.70 %; transfer-stimuli trials mean = 93.10 %, SD = 1.81 %) showed above chance-level performance (all ps < .001) with no difference between known- and transfer-stimuli trials (all *p*s > .250). There was no difference between stimuli of class X and Y in both known-(t(3) = 0.79, *p* > .250, Cohen’s d = 0.37) and transfer-stimuli trials (t(3) = 0.53, *p* = > .250, Cohen’s d = 0.33).

For the four animals that were tested in non-reinforced transfer conditions, we found that the CC classifier could predict the presented stimulus class above chance-level in known-stimuli trials (P580: 84.16 %, P592: 74.16 %, P593: 66.80 %, P599: 78.20 %, all *p*s < .001, see supplementary figure 11A).

CC classifier accuracy was also above chance-level in transfer-stimuli trials (P580: 84.48 %, P592: 71.28 %, P593: 61.60 %, P599: 76.68 %, all *p*s < .001, see supplementary figure 11B). Classifier accuracy did not differ in transfer- compared to known-stimuli trials except for pigeon P593. Here, classifier accuracy dropped significantly in transfer-stimuli trials (t(9) = 3.77, *p* = .004, Cohen’s d = 1.20).

For pigeons P592 and P593, we could also analyze known-stimuli error trials as there were sufficient error events to warrant further analysis. For pigeon P592, we found a significantly lower than chance classification accuracy when using pecks from correct trials as class predictors and error pecks as test events (35.24 %, t(9) = 9.33, *p* < .001, Cohen’s d = 2.96, see supplementary figure 11C). This indicated that P592 pecked on features relevant for stimulus class X even though a class Y stimulus was presented in error trials and vice versa (see supplementary data for the pecking responses of individual pigeons). There was no difference from chance in the CE classification (t(9) = 1.76, *p* = .112, Cohen’s d = 0.55) for pigeon P593 indicating that there was no association between pecking locations of correct and error trials. For neither pigeon, transfer-stimuli trials could be analyzed in the CE classification due to low number of error trials.

There were sufficient errors in known-stimuli trials for pigeons P592 and P593 to perform an EE classification in which the classifier was both trained and tested on pecking events during erroneous trials. For both pigeons, we found no evidence that pecking in error trials could be predict the presented stimulus class in other error trials (P592: 56.56 %, P593: 52.92 %, both ps >. 085). There were not sufficient error events in transfer-stimuli trials preventing further analysis.

