## Supplementary figures for "Digital embryos – A novel technical approach to investigate perceptual categorization in pigeons (*Columba livia*) using machine learning"

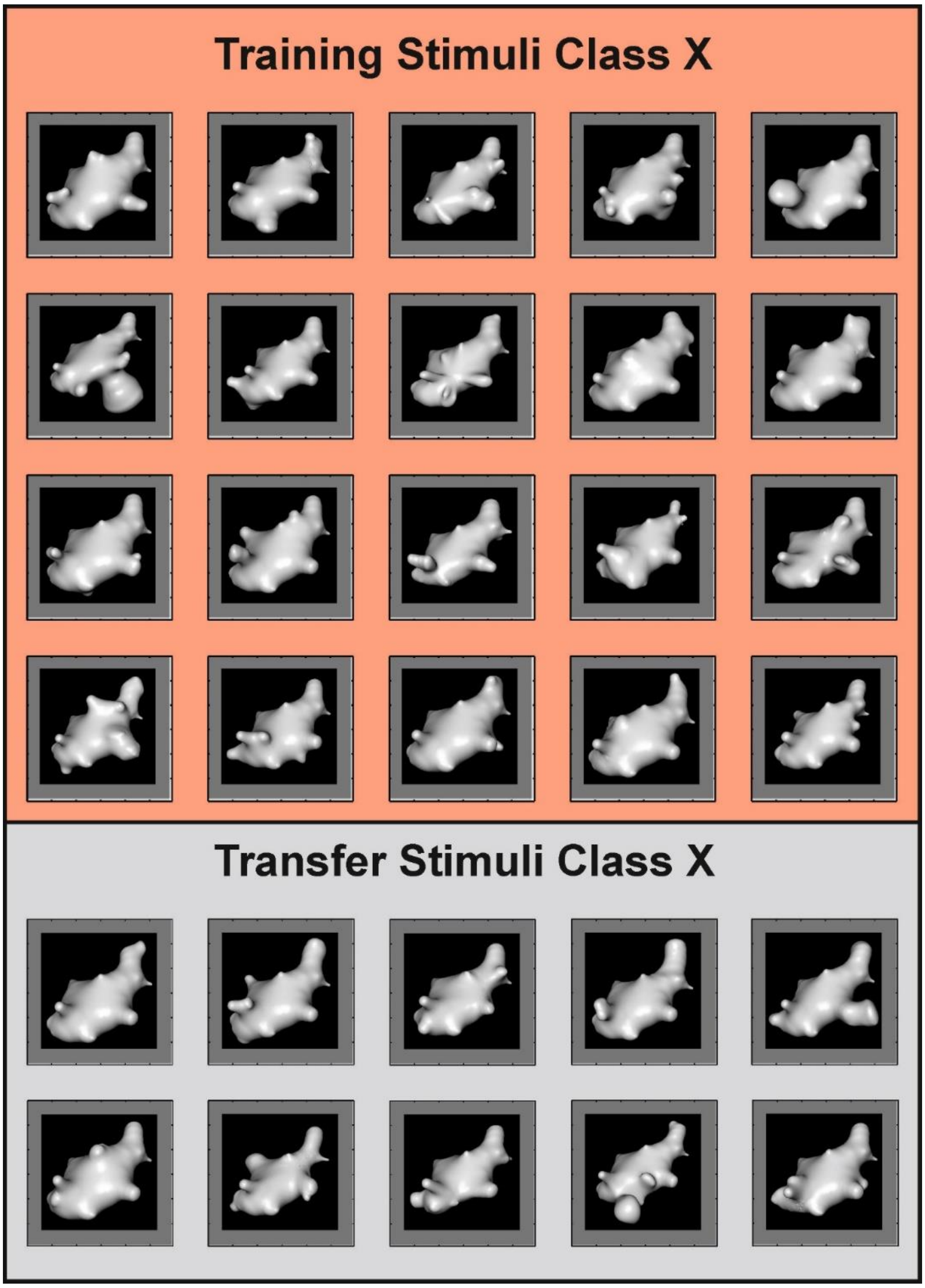


**Supplementary figure 1. Example training and transfer stimuli for class X from one session.**

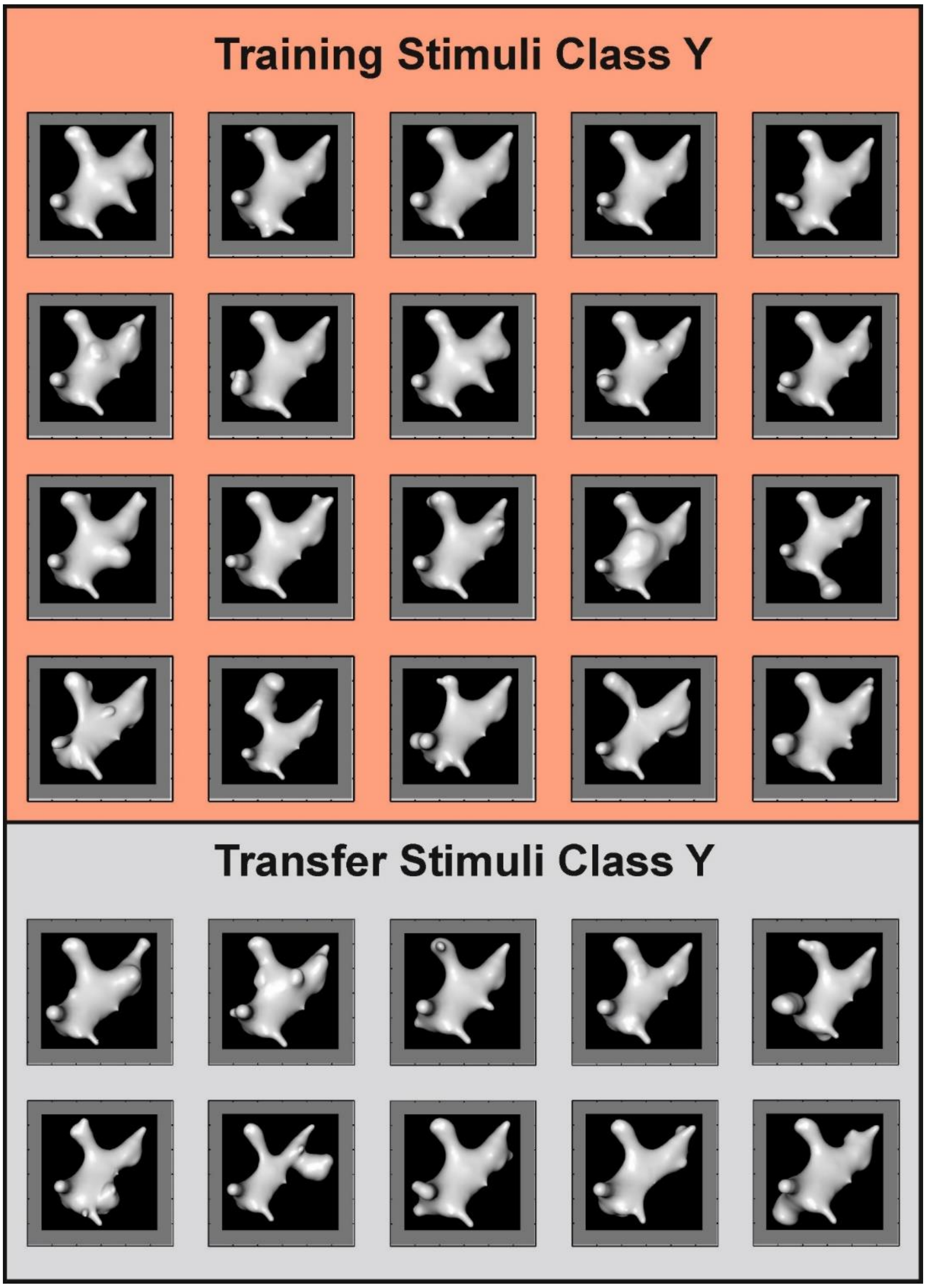


**Supplementary figure 2. Example training and transfer stimuli for class Y from one session.**

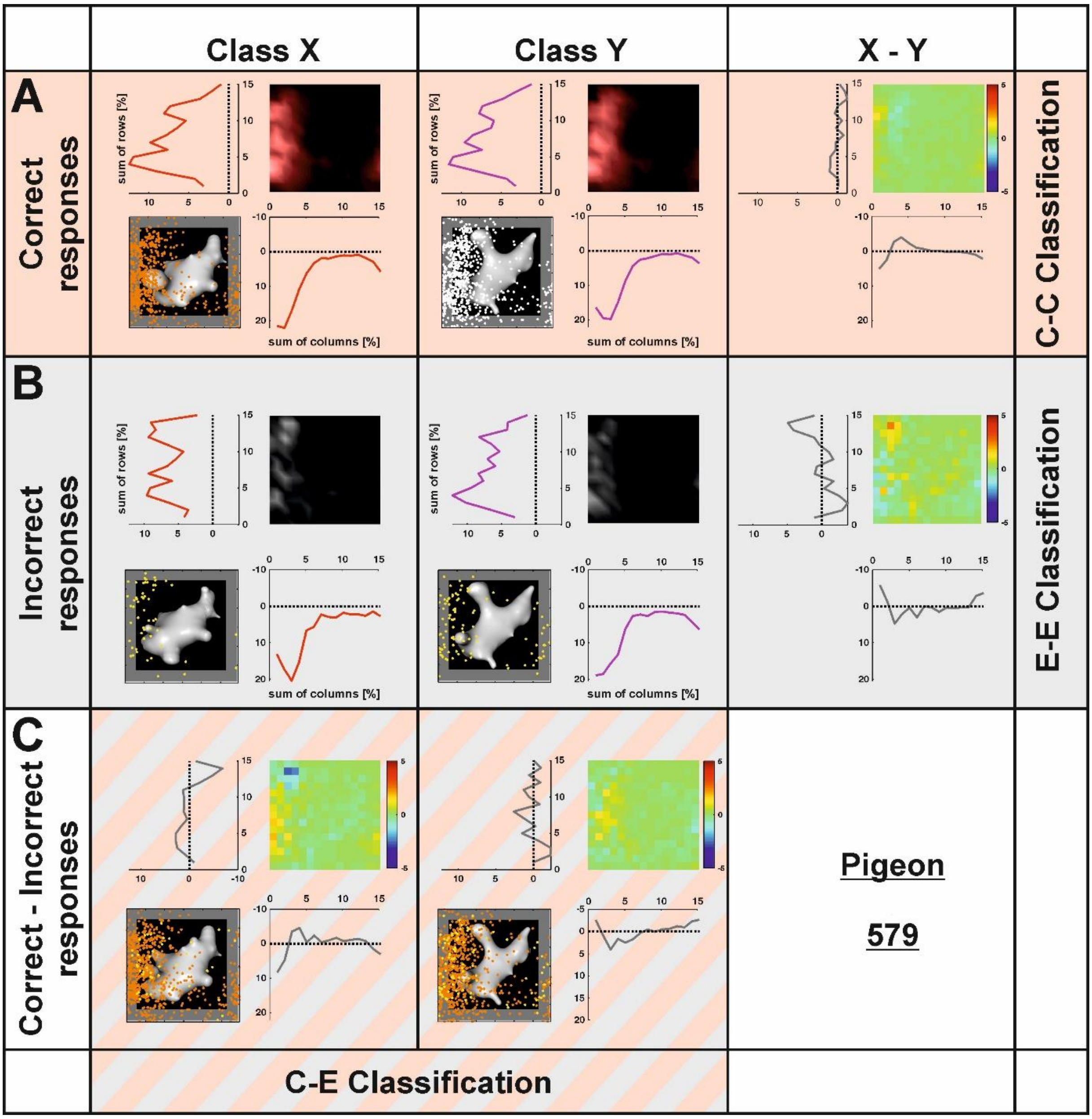


**Supplementary figure 3. Heatmap analysis for pigeon 579.**

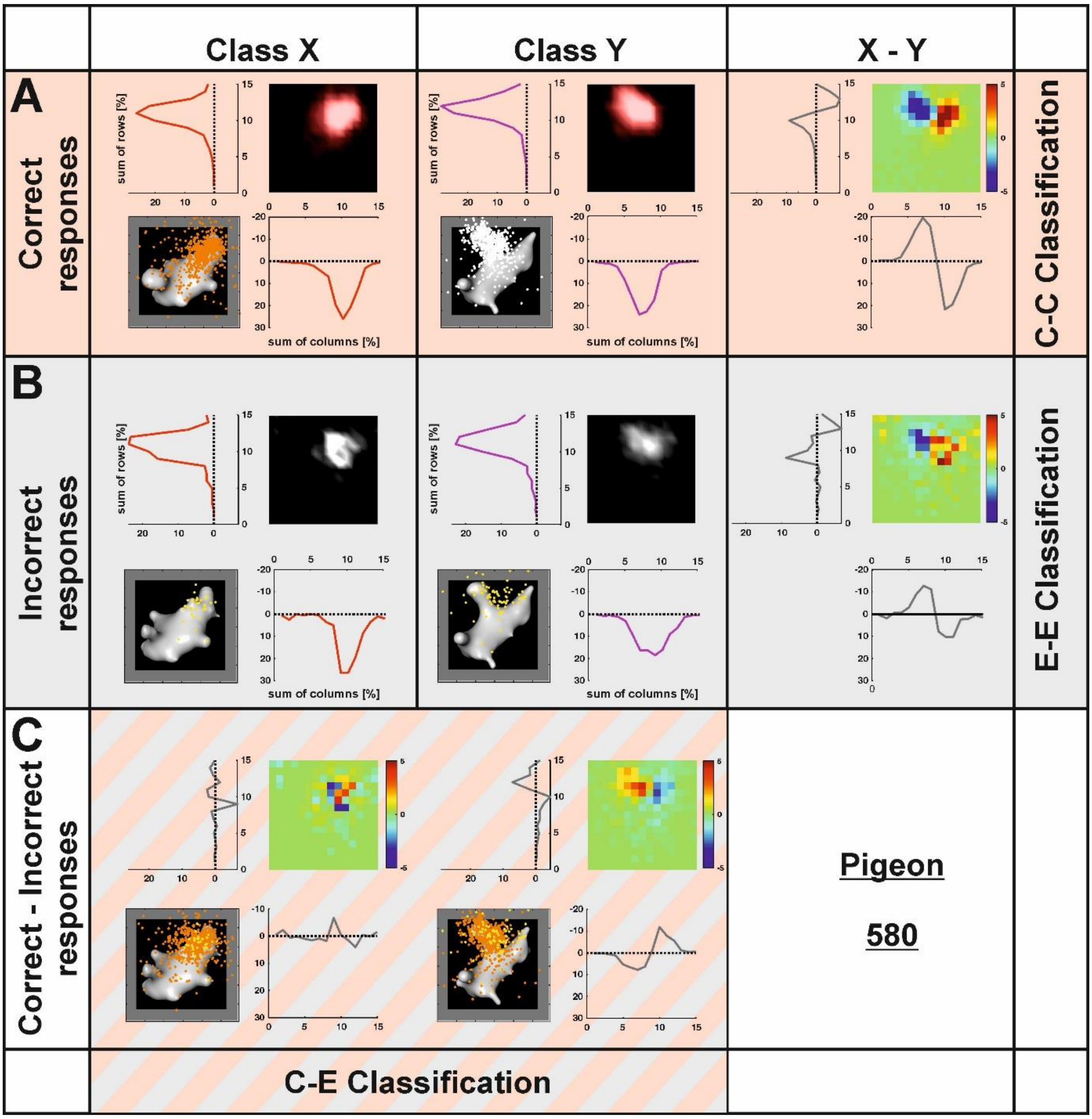


**Supplementary figure 4. Heatmap analysis for pigeon 580.**

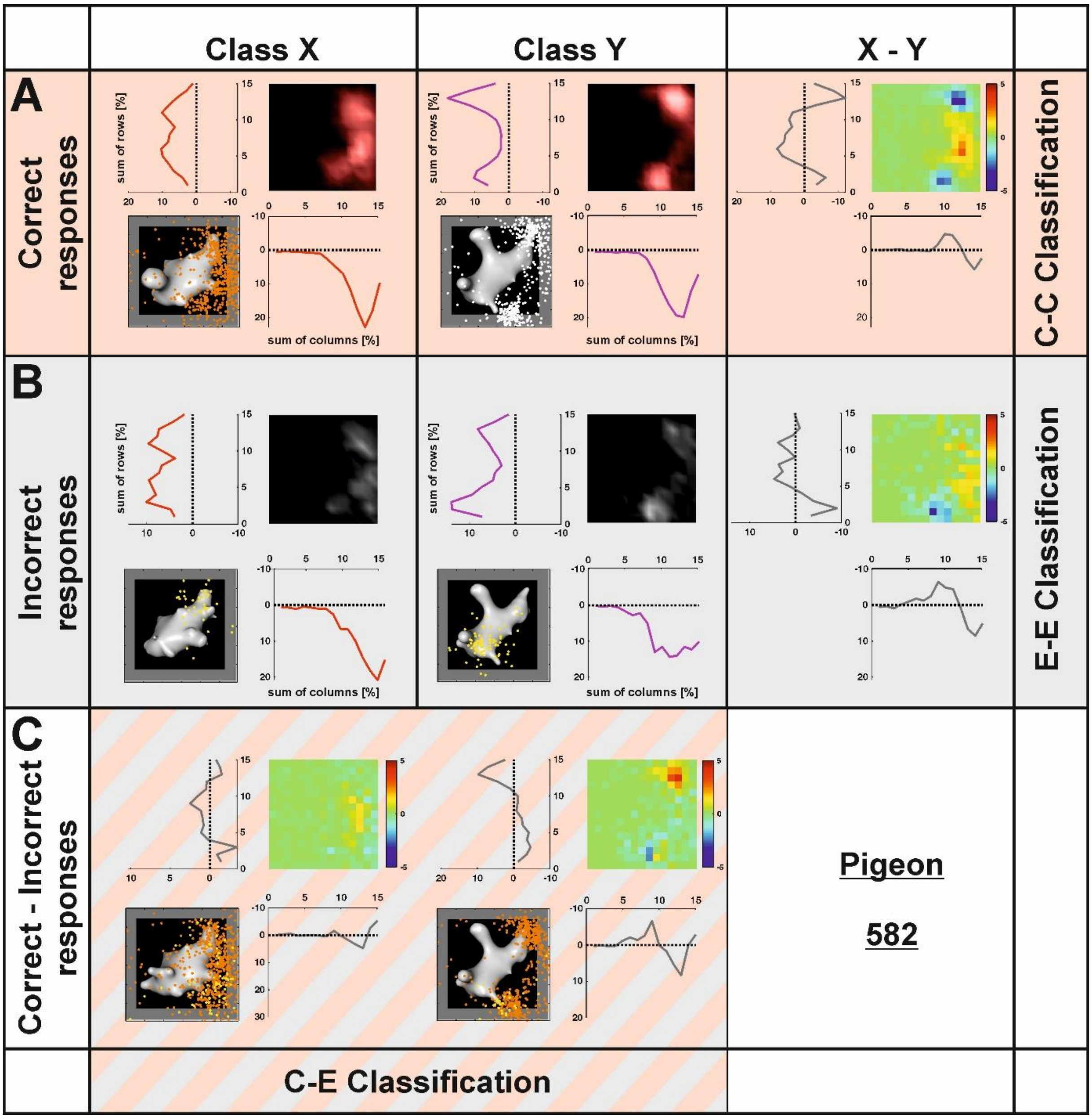


**Supplementary figure 5. Heatmap analysis for pigeon 582.**

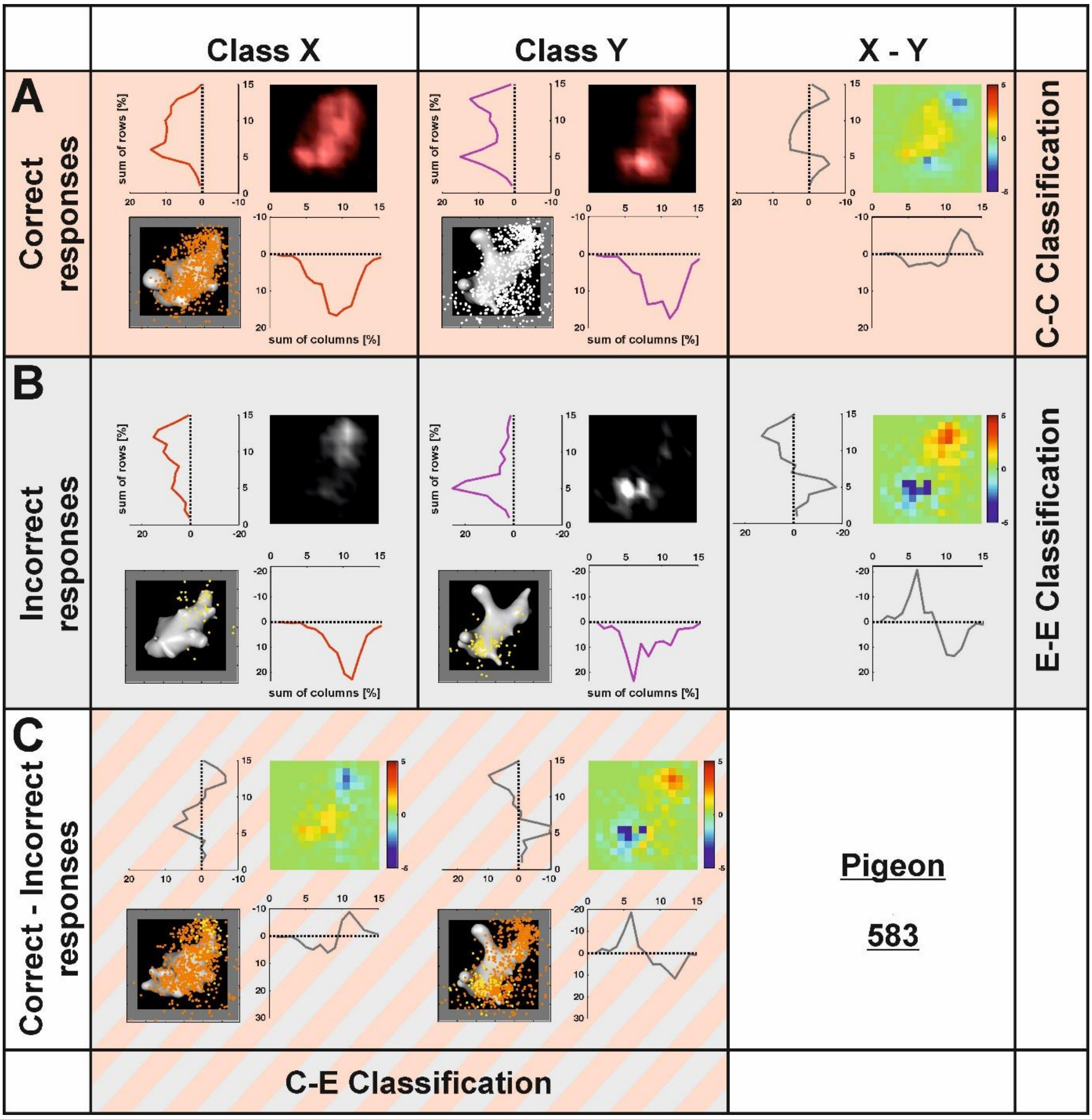


**Supplementary figure 6. Heatmap analysis for pigeon 583.**

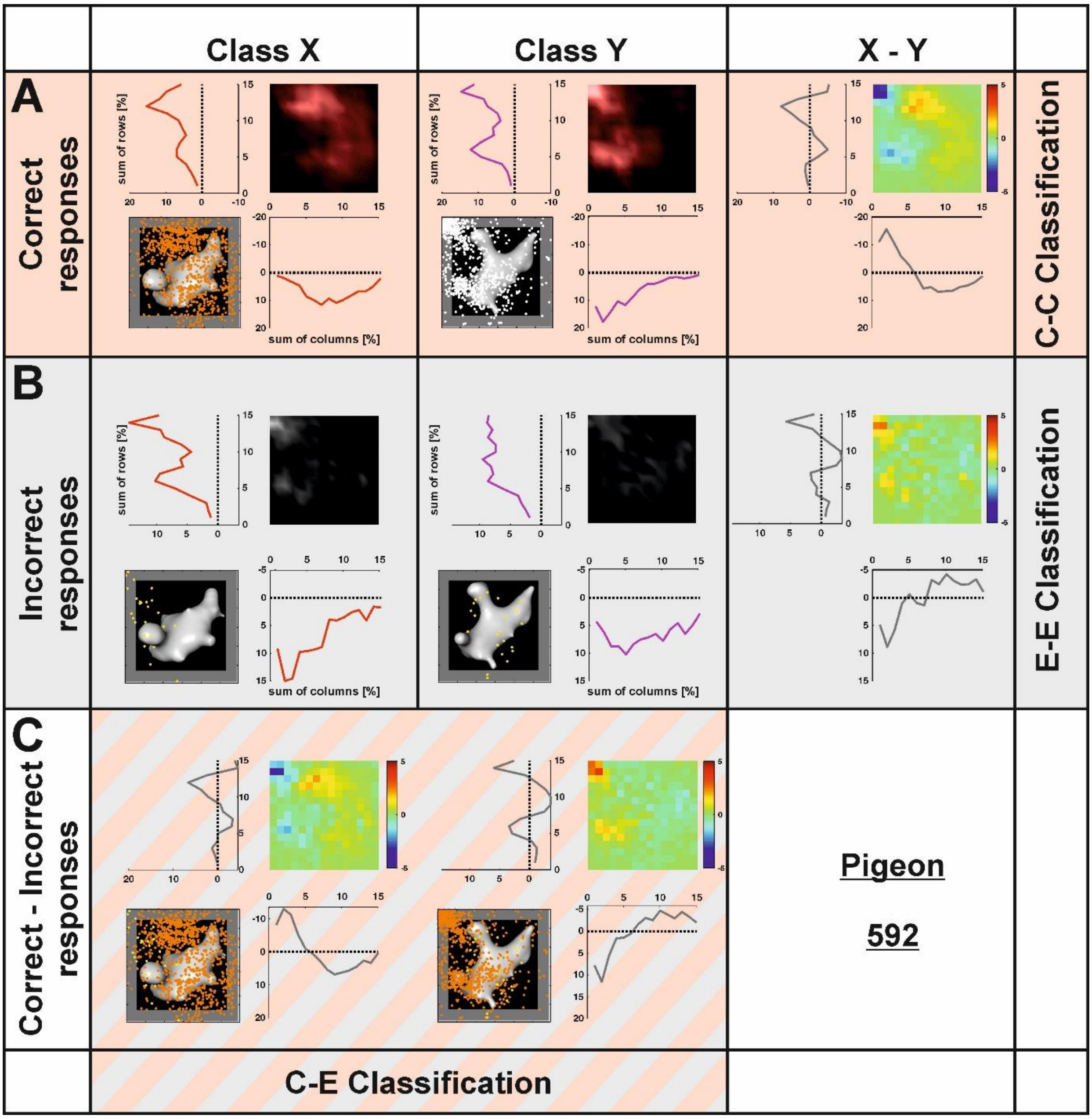


**Supplementary figure 7. Heatmap analysis for pigeon 592.**

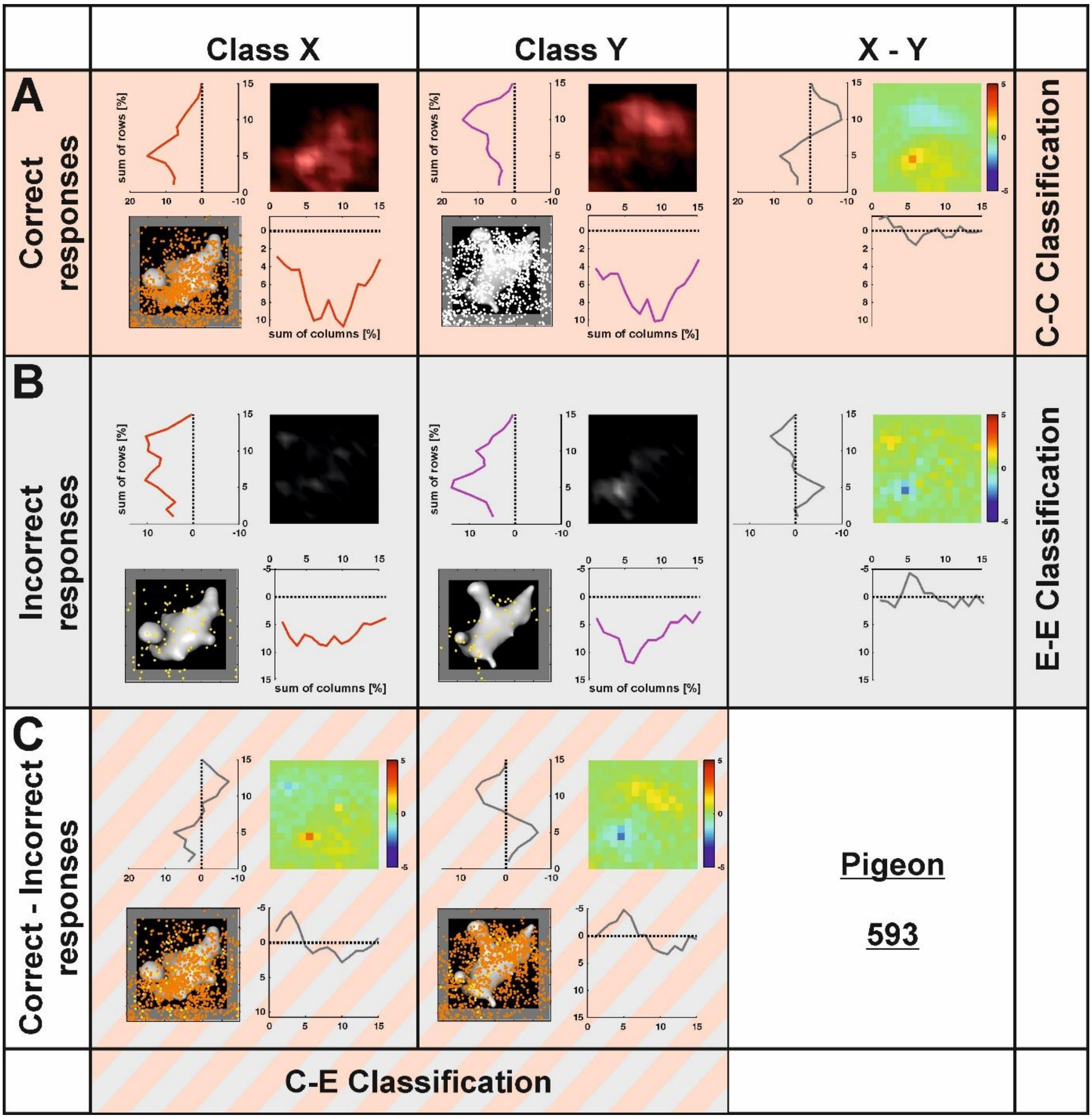


**Supplementary figure 8. Heatmap analysis for pigeon 593.**

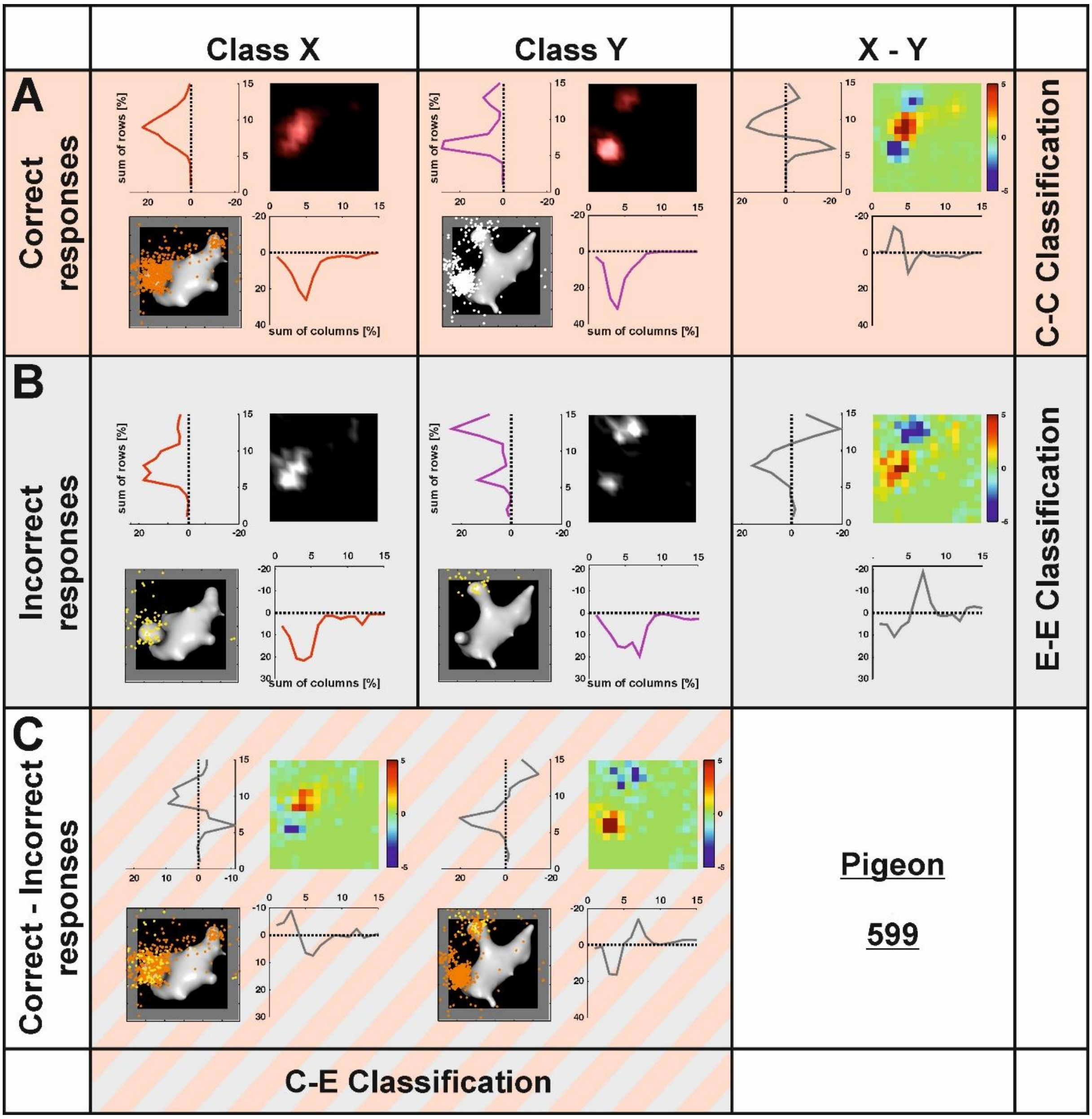


**Supplementary figure 9. Heatmap analysis for pigeon 599.**

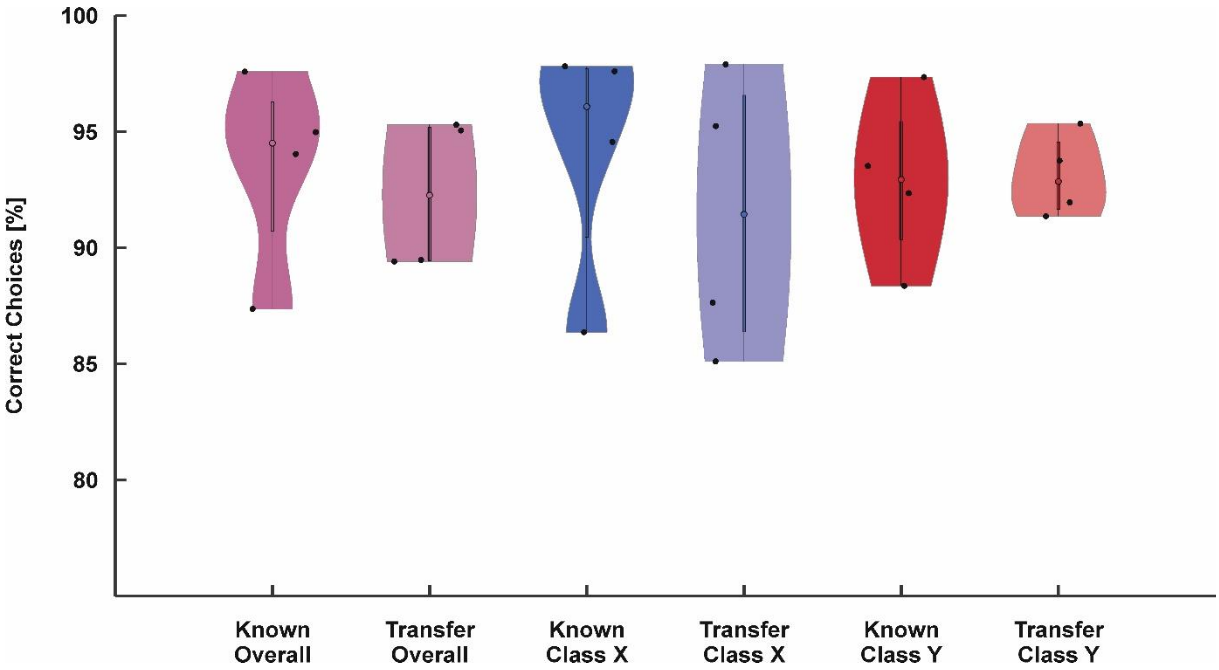


**Supplemetary figure 10. Behavioral results**

Violin plot of behavioral performance for known and transfer trials overall and broken down by embryo class. Digital embryos could be categorized in each experimental condition in non-reinforced circumstances. There was no difference between the performance to the known embryos and the transfer to new instances of embryo classes X and Y in any of both conditions.

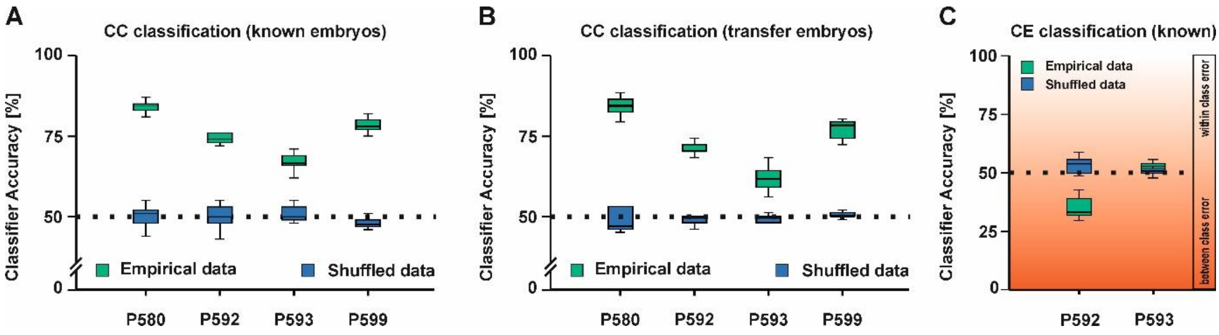


**Supplemetary figure 11. CC and CE classification results of the non-reinforced transfer**

Digital embryos could be classified for each animal tested (**A** shows the classifier response for the known stimuli and **B** depicts the classifier results for the transfer stimuli). **C** shows the classifier response for the CE classification. Dark red colors indicate a confusion between the categories X and Y. Light red colors indicate confusion within a given category in the CE classification.
